# Cytokinin regulates the gravitropic setpoint angle of seminal roots in barley and wheat

**DOI:** 10.64898/2026.10.09.757912

**Authors:** Atish Bansod, Sophie Zoe Farkas, Miriam Schreiber, Tracy A. Valentine, Aleš Pěnčík, Ondřej Novák, Petra Amakorová, Jürgen Kleine-Vehn, Rahul Bhosale, Gwendolyn K. Kirschner

## Abstract

Plant root growth angle determines the spatial distribution of roots in soil and thereby influences access to water and nutrients when these resources are heterogeneously distributed. In barley, multiple seminal roots emerge at distinct gravitropic setpoint angles (GSAs) and they reliably re-establish their characteristic growth angles following reorientation, indicating robust intrinsic control. However, the molecular basis underlying GSA variation among roots within a single root system remains unknown. Here, we combined individual root type-specific transcriptomics with laser-capture microdissection to profile gravity-responsive zones (root cap, meristem, and elongation zone). We identified two *CYTOKININ OXIDASE/DEHYDROGENASE* (*CKX*) genes whose expression in the root cap was associated with root-specific GSA. Functional analysis showed that *ckx* mutants, which show higher cytokinin levels, exhibit significantly steeper seminal root GSAs, implicating cytokinin catabolism in root angle regulation. Consistent with this, exogenous cytokinin treatment induced a dose-dependent steepening in root angle in barley. Analyses of gravitropic curvature after gravistimulation further showed that *ckx* mutants exhibit altered gravitropic responses, indicating that *CKX*-mediated regulation of the root angle depends on gravity-responsive pathways. Notably, GSA-associated *CKX* expression patterns and cytokinin-dependent root angle responses were also observed in wheat, suggesting a conservation of this mechanism across cereals. Together, these findings identify cytokinin as a key regulator of root specific setpoint angles within a root system architecture, with potential implications for optimizing crop resource acquisition and yield.

## Introduction

Plant root angle determines rooting depth and thereby the ability of the root system to access water and nutrients located in different soil layers (1). Root systems with steeper root angles place more roots in deeper soil layers, where deep-soil water and mobile nutrients are present. In contrast, root systems with shallower root angles have more roots in the topsoil, where relatively immobile nutrients such as phosphate accumulate. Accordingly, plants with steeper root systems are generally considered better adapted to drought conditions, as shown for the *DEEPER ROOTING1* allele in rice (2). Near-isogenic lines carrying *DRO1* show steeper root angles and maintain high yield performance under drought conditions, presumably because they can access deep-soil water better than plants with shallower root systems.

The root angle is set in relation to gravity. This gravitropic-setpoint angle (GSA) is defined as the angle at which a plant organ is maintained as a result of gravitropism (3). In roots, gravity perception, signalling and response takes place at the root tip. In response to a change in direction of gravity such as after rotation, amyloplasts in the root cap change position and settle in the cells towards the new centre of gravity. This gravistimulation triggers a Mitogen-Activated Protein Kinases (MAPK) cascade that phosphorylates LAZY (LZY) proteins and promotes their enrichment on the surface of amyloplasts (4). LZY proteins subsequently translocate to the plasma membrane adjacent to the amyloplasts and thereby transmitting their positional information to the plasma membrane (4, 5). At the plasma membrane, the polarly localised LZYs recruit RCC1-like domain (RLD) proteins, which promote the re-localization of the auxin transporters PINFORMED (PINs) and modulate auxin flow (5). This process generates a lateral auxin gradient with auxin accumulatingin lateral root cap and epidermis cells on the lower side of the root (6), where AUXIN RESPONSE FACTOR 19 (ARF19) functions as a transient enhancer of auxin signalling, reinforcing differential growth responses required for root bending response (7). Enhanced auxin levels subsequently induce apoplastic pH changes in a Ca^2+^-dependent signaling pathway and accumulation of reactive oxygen species (ROS), leading to differential cell expansion in the elongation zone (8, 9). This directs the root growth towards to the new center of gravity.

This mechanism has been extensively studied in Arabidopsis primary roots that have a vertical GSA. In the case of non-vertical GSAs like in Arabidopsis lateral roots, it has been proposed that the non-vertical growth is initiated by differential distribution of auxin and growth repression in the lower flank of young lateral roots (10). Auxin-dependent gravitropic growth is counterbalanced by asymmetric cytokinin signaling at the upper root flank, which reduces differential growth and ultimately bending to gravity (11, 12). In later growth stages, the lateral roots maintain their GSA through the balance of downward gravitropic and upward antigravitropic (AGO) auxin fluxes generated in the columella and mediated by PIN3 and PIN7 (13). Hence, a non-vertical GSA results from a balance between gravitropic and AGO forces.

In cereals, root systems consists of more root types that show non-vertical GSA, including embryonic seminal roots and shoot-borne crown and brace roots (14). Several mechanisms that regulate gravitropic growth in these roots have been identified. In rice and maize, ethylene positively influences root gravitropism via stimulating MAO HU ZI10/TRYPTOPHAN AMINO-TRANSFERASE2-mediated auxin biosynthesis (15). Similar to ethylene, abscisic acid governs root gravitropism by inducing auxin synthesis (16). Furthermore, rice ARFs promote a positive gravitropic response by activating *Increased Leaf Angle1* expression in epidermal cells on the lower side of the root. This increases cell wall thickness and thereby inhibiting cell elongation, leading to gravitropic bending (17). Additionally, key players in the AGO mechanism have been identified in wheat (*Triticum aestivum*) and barley (*Hordeum vulgare*) (18, 19). Mutations in *ENHANCED GRAVITROPISM1* (*EGT1*) and *EGT2* result in hypergravitropic root growth with all root types showing a near vertical GSA. Both genes function in parallel pathways and are involved in cell wall regulation, suggesting that they act in the AGO pathway by stiffening cell walls and counteract the auxin-dependent gravitropic mechanism.In wheat, these pathways modulate a conserved gravitropic response to generate root class-specific GSAs (20).

Together, these mechanisms explain how roots of the same root type establish and maintain their specific GSA. For example, in Arabidopsis, lateral roots of similar developmental age exhibit highly consistent GSAs. Interestingly, however, some root types maintain different GSAs despite sharing the same genetic background and developmental stage. A striking example is provided by barley seminal roots (Fig. 1A). Barley seminal roots not only maintain their characteristic GSAs during elongation, but also return to these angles after reoreintation (18) (Suppl. Fig. 1C and D), suggesting intrinsic differences among individual roots in their gravitropic and AGO responses. To investigate the basis of these differences, we studied the transcriptome of the root cap, the meristem and elongation zone in barley seminal roots with different GSAs and identified key genes that were differentially expressed between roots with different growth angles. Based on their proposed role for GSA regulation in Arabidopsis lateral roots, we focused on *CYTOKININ OXIDASE/DEHYDROGENASE* (*CKX*) genes. We found that increasing cytokinin levels in barley seminal roots through cytokinin application or mutations in *CKX* genes, induced steeper root growth. Moreover, this response was conserved in the embryonic root system of wheat. We therefore propose that cytokinin homeostasis contributes to the fine-tuning of gravity responsiveness among individual roots and thereby regulates their GSA.

**Figure 1:**
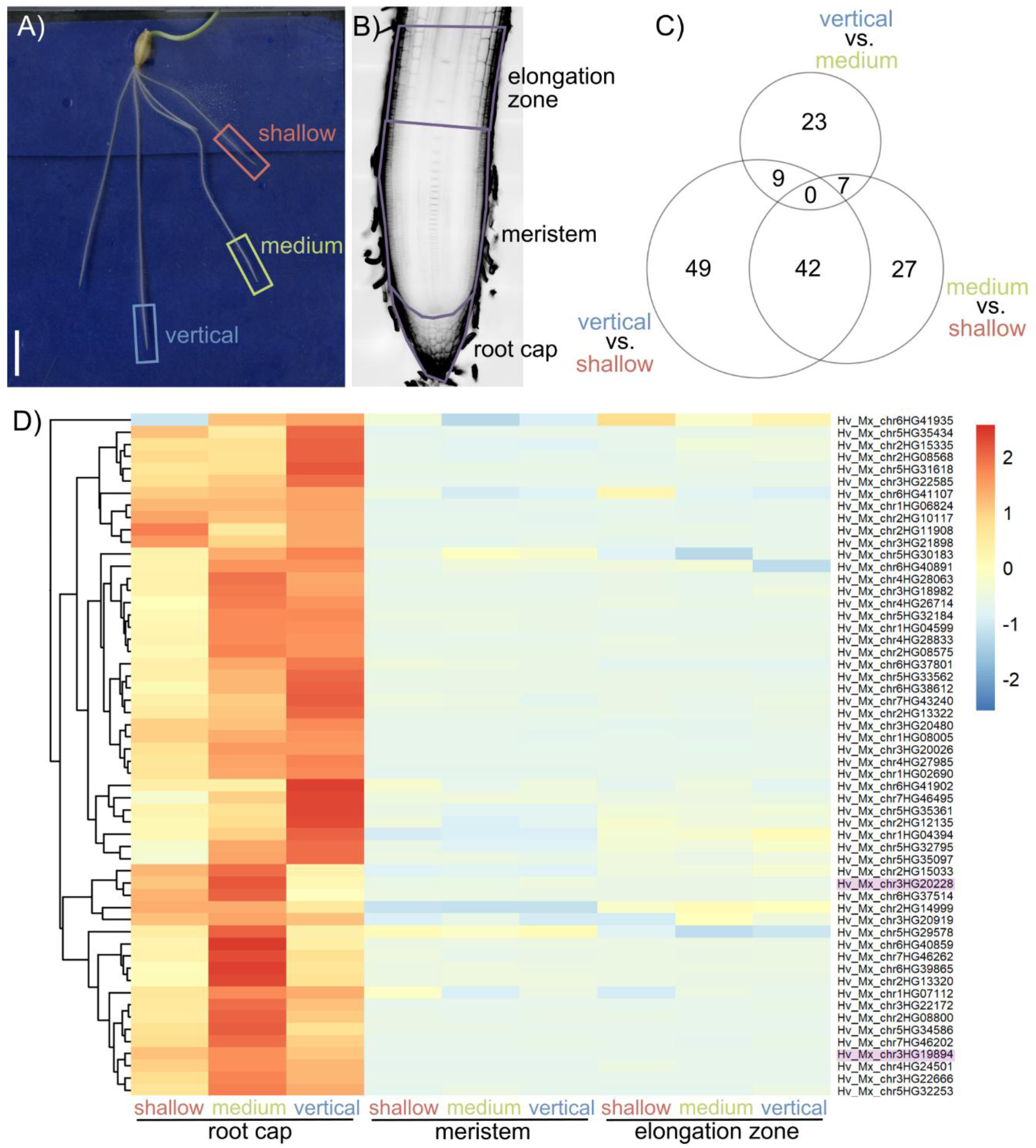
Barley seminal roots with different gravitropic set-point angles have different transcriptomic signatures. A) Roots for RNA-seq were sampled from seedlings of the barley cv. Morex 4 days after germination; vertical root had an angle of 85-90° from horizontal; medium angle roots had an angle of 60-70° from horizontal; shallow roots had an angle of 40-50° from horizontal; scale bar 1 cm. B) Root cap, root meristem and elongation zone were sampled by laser capture microdissection. C) Venn diagrams showing differentially expressed genes (DEGs) in the respective comparisons in the root cap. D) Heatmap of differentially expressed genes, showing genes predominantly expressed in the root cap (complete heatmap can be found in Supp. Fig. 3); *HvCKXs* are highlighted in purple; colours show z-score according to colour scale.

## Results

### Different gravitropic set-point angles among seminal roots are associated with distinct transcriptomic signatures

Barley seminal roots are established in the embryo and emerge during the first days after germination. Consequently, they are of similar age and developmental stage. However, they exhibit distinct GSAs (Fig. 1A). Analysis of starch content and distribution did not reveal any differences in the amyloplasts that serve as statoliths in the root cap among roots with different GSAs (Suppl. Fig. 1A, B). Interestingly, following gravistimulation by a 90° rotation, all roots gradually returned to the GSA they had before rotation (Suppl. Fig. 1C, D). Roots whose growth angle remained unchanged by the rotation, for example, roots growing at 45° relative to the horizontal before and after the rotation, continued growing in the same direction. In contrast, roots that exhibited a growth angle of 45° before rotation but were placed at a −45° angle after rotation (i.e., became shallower than horizontal) progressively redirected their growth back toward 45° over time.

To test whether this intrinsic GSA memory was associated with distinct transcriptomic signature, we generated and analyzed the transcriptome of roots with different GSAs: a vertical GSA (85-90° from horizontal); a medium GSA (60-70° from horizontal); and a shallow GSA (40-50° from horizontal) (Fig. 1A). To capture zone-specific differences, we used laser-capture microdissection (LCM) to isolate RNA from gravity-responsive zones, namely the root cap, the meristem and the elongation zone (Fig. 1B, Suppl. Fig. 2A). Through within-zone comparisons of roots exhibiting different GSAs, we identified 152 differentially expressed genes (DEGs) (adjusted *P* < 0.05 and abs (log2 fold change) ≥1, the vast majority of which were detected in the root cap (149 DEGs). In contrast, only 3 DEGs were identified in the meristem, and none were detected in the elongation zone (Fig. 1C, Suppl. Table 2). Several DEGs were shared among multiple comparisons of root cap samples from roots with different GSAs (Fig. 1C, D). The relatively low number of DEGs highlights the high degree of similarity among the sampled roots with respect to age and developmental stage, indicating that they differ primarily in their GSAs. This observation suggests that the identified DEGs may play important roles in the establishment and maintenance of GSA.

### Cytokinin is involved in the regulation of the gravitropic set-point angle in barley and wheat seminal roots

Within the 152 DEGs, we did not identify any enriched biological processes, molecular functions, or cellular components through gene ontology enrichment analysis. Therefore, we focused on candidate genes based on findings from previous studies in other plant species. In Arabidopsis, a genome-wide association study (GWAS) approach and mutant phenotyping showed that cytokinin metabolism mediated by CKX2 is involved in the regulation of lateral root GSA (11). Among the DEGs in the root cap, we identified two genes encoding cytokinin oxidase/dehydrogenases (Hv_Mx_chr3HG19894 and Hv_Mx_chr3HG20228), designated before as *HvCKX1* and *HvCKX5*, respectively (21) (Suppl. Fig. 3). Out of all CKX encoding genes in barley, Hv_Mx_chr1HG04074 (previously designated as *HvCKX3* (22)) and Hv_Mx_chr7HG48516 were also expressed in the examined zones (Suppl. Fig. 4). These findings suggests that cytokinin may play a role in the regulation of GSA in barley seminal roots, similar to its reported function in Arabidopsis lateral roots (11, 12).

To investigate the influence of cytokinin levels on GSA, we treated barley seedlings with the synthetic cytokinin 6-benzylaminopurine (6-BA). Cytokinin treatment resulted in a narrower root system, with increasing cytokinin concentrations causing a dose-dependent reduction in root angle (Fig. 2A, B), suggesting that cytokinin levels influence GSA in barley seminal roots. Regulation of seminal root angle is often conserved among cereals at both the genetic (18, 19) and hormonal levels (15). Therefore, we examined whether cytokinin-mediated regulation of seminal root angle is conserved among cereal species. Cytokinin treatment of wheat seedlings similarly altered root angle, resulting in a steeper root system at higher cytokinin concentrations (Fig. 2C, D). These findings indicate that cytokinin-dependent regulation of seminal root angle is conserved between barley and wheat.

**Figure 2:**
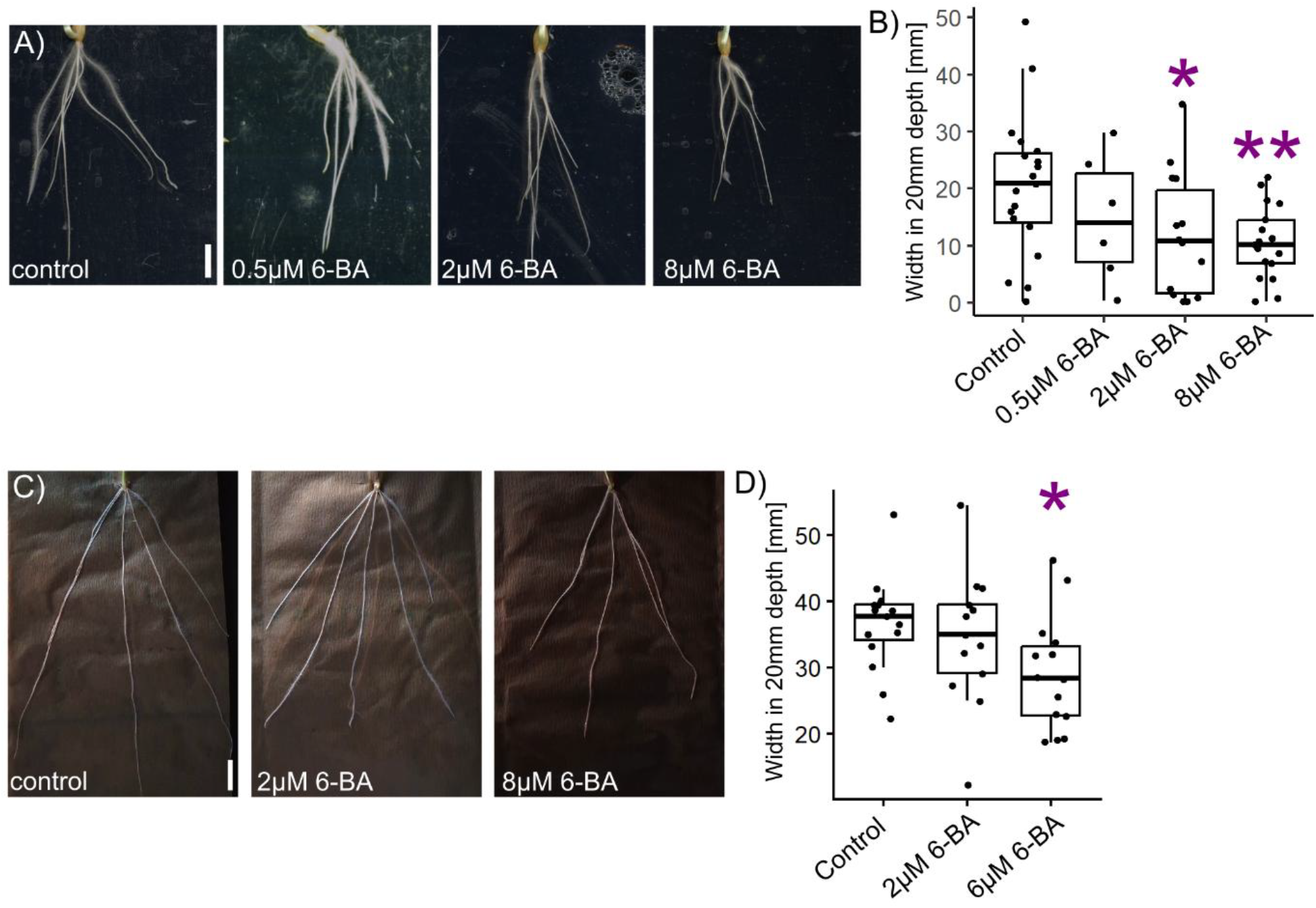
Cytokinin treatment steepens the root angle in a dose-dependent manner. A) Barley cv. Morex grown on media containing no (control), 0.5µM, 2µM or 8 µM 6-BA 6-Benzylaminopurine (6-BA) for 5 days after germination; scale bar 1 cm. B) Root system width of barley cv. Morex seedlings grown on media containing 6-BA 20 mm downwards of the seed; n = 19 plants (control), 6 (0.5 µM 6-BA), 14 (2 µM 6-BA), 17 (8 µM 6-BA); experiment was performed twice. C) Wheat cv. Cadenza grown on water containing no (control), 2 µM or 6 µM 6-BA for 6 days after germination; scale bar 2 cm. D) Root system width of wheat cv. Cadenza seedlings grown on media containing 6-BA ; n = 15 plants (control), 13 (2 µM 6-BA), 14 (6 µM 6-BA); asterisks mark samples that are significantly different from WT (Student’s t-test, * = p<0.05, ** = p<0.01).

### Comparison of wheat and barley transcriptomes highlights a role for cytokinin oxidase-encoding genes in the root gravitropic set-point angle

Since the root angle response to cytokinin application was conserved between wheat and barley, we analysed the expression of the wheat orthologues of *HvCKX1, HvCKX3* and *HvCK5* in root tips of wheat seminal roots (Fig. 3A) (20). In wheat, the embryonic seminal roots emerge in a defined temporal sequence, with each root class exhibiting a distinct GSA. The primary root emerges first after germination and grows nearly vertically. This is followed by the first pair of seminal roots, which adopt a shallower GSA, and finally by the second pair of seminal roots, which emerge at an even shallower growth angle (Fig. 3B) (23).

**Figure 3:**
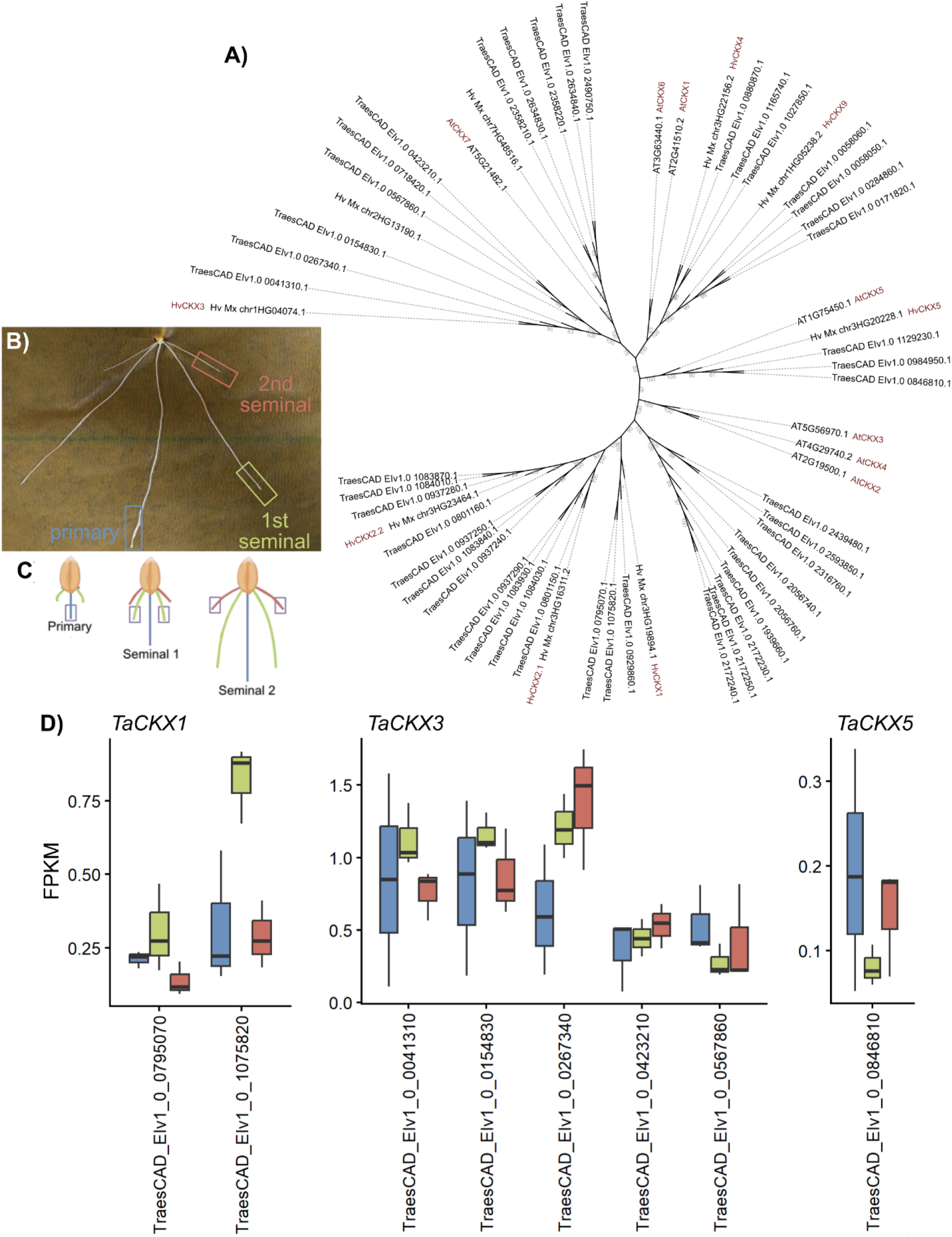
*CYTOKININ OXIDASE/DEHYDROGENASE* (*CKX*) orthologues in wheat are expressed in a root-class specific manner. A)Phylogenetic tree of CKXs in Arabidopsis, wheat and barley. B)The embryonic root system in wheat exhibits distinct gravitropic set-point angles (GSAs); the primary root grows vertically, seminal root pair 1 in a medium-shallow angle and seminal root pair 2 more shallow. C)Root tips for RNA-seq were sampled to reflect similar developmental stages and age of the different embryonic roots. D) Expression of wheat *CKXs* in 1 cm long roots of different root classes, primary root (red), first pair of seminal roots (yellow) and second pair of seminal roots (red).

Similar to *HvCKX1* and *HvCKX5* in barley, their wheat orthologues were expressed in a root class-specific manner, exhibiting either the highest or lowest expression levels in root classes with medium shallow angle, respectively (Fig. 3C, D, Suppl. Table 6). In addition, several wheat orthologues of *HvCKX3* showed expression patterns that correlated with root GSA. These findings suggest that *CKX* genes are associated with root angle variation among seminal root classes. Furthermore, the distinct expression patterns of individual CKX family members indicate that they may perform specialised functions in the establishment and maintenance of root class specific GSAs.

### *CYTOKININ OXIDASE/DEHYDROGENASE* genes are involved in the gravitropic set-point angle regulation in barley seminal roots

To investigate the role of *HvCKX*s genes in root angle regulation, we utilised previously characterised mutants in *HvCKX1* and *HvCKX3* (22). Seminal roots of *Hvckx1-1, Hvckx1-2* and *Hvckx3* showed signficantly steeper root GSAs compared to the wild type (WT) (Fig. 4A, B). A similar phenotype was observed in plants grown in soil and visualised using X-ray micro-computed tomography (μCT) (Fig. 4C, D, Suppl. Mov. 1-4).

**Figure 4:**
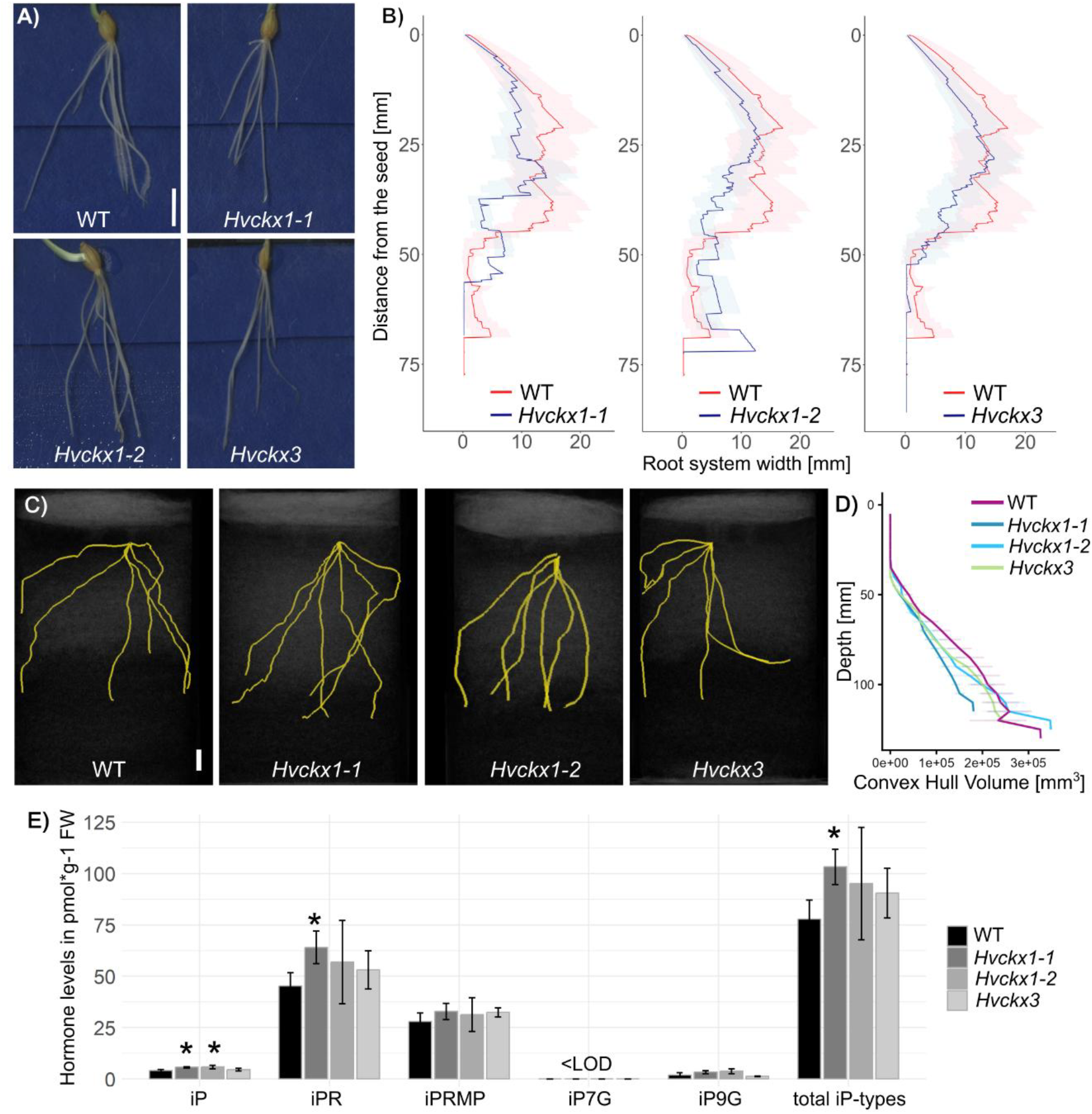
*CYTOKININ OXIDASES DEHYDROGENASES* (*CKXs*) regulate the gravitropic set-point angle (GSA) in barley. A*) Hvckx* mutants have a steeper seminal root angle than wild type (WT) Golden Promise; seedlings 4 days after germination; scale bar 1 cm. B) Root system width plotted by distance to the seed; lines show mean of root system width, the error bars show the standard deviation; n = 17 plants (WT), 10 (*Hvckx1-1*), 13 (*Hvckx1-2*), 22 (*Hvckx3*). C) X-ray microcomputed tomography (μCT) images of WT, *Hvckx1-1, Hvckx1-2* and *Hvckx3* grown for xxx days after germination in soil columns; scale bar 1 cm. D) Convex hull volume of WT and *Hvckx* mutants quantified from x-ray μCT; n = 8 plants (WT), 6 (*Hvckx1-1*), 8 (*Hvckx1-2*), 8 (*Hvckx3*). E) Levels of iP-type cytokinins in WT, *Hvckx1-1, Hvckx1-2* and *Hvckx3* root tips; n = 4 plants (WT), 5 (*Hvckx1-1*), 4 (*Hvckx1-2*), 4 (*Hvckx3*) 4 days after germination; mean and standard deviation are shown; asterisks mark samples that are significantly different from WT (Student’s t-test, p<0.05).

Following gravistimulation by 90º reorientation, *Hvckx1-1* and *Hvckx1-2* displayed steeper root bending angles than WT 24 h after the rotation, whereas *Hvckx3* showed a response comparable to that of the WT (Supp. Fig. 5A, B). Quantification of cytokinin levels revealed that *Hvckx1-1* and *Hvckx1-2* contained higher levels of isopentenyladenine base (iP)-type cytokinins than the WT, whereas *Hvckx3* contained lower levels of cis-zeatin (cZ)-type cytokinins (Fig. 4E, Supp. Table 5).

Together with the steeper root angle phenotype observed in *Hvckx1* and *Hvckx3* mutants, these results suggest that the homeostasis of different types of cytokinin is important for maintaining GSA in barley seminal roots. Furthermore, the altered gravitropic responses of the *Hvckx* mutants indicate that cytokinin metabolism contributes to root angle regulation, at least in part, through modulation of gravity responsive growth processes.

## Discussion

The term GSA was first introduced by Digby and Firn in 1995 to describe the angle at which a plant organ is maintained relative to the gravity vector (3). Interestingly, seminal roots in barley arising from the same plant can exhibit different GSAs, even though they develop at a similar time and share the same genetic background, environment and exposure to gravity. Following a misplacement from the original gravity vector by reorientation, these seminal roots adjust their growth direction and return to their characteristic set angle (Fig. 1, Supp. Fig, 1). A similar process is observed in wheat, where the primary root and the successive seminal root pairs emerge and maintain different GSAs (20, 23). To date, the genetic factors underlying the maintenance of different GSAs among roots of the same type within a single plant remain largely unknown. Likewise, it is unclear how individual roots modulate their responsiveness to the same directional cue to establish distinct growth trajectories. Here, we compared the transcriptome of barley seminal roots exhibiting different GSAs, ranging from steep to shallow. Roots were separated into three distinct root zones, namely the root cap, meristem, and elongation zone, using LCM, because gravity perception, signal transduction and differential cell elongation occur primarily within these zones. Interestingly, most DEGs were identified in the root cap (Fig. 1). Although LCM enables precise isolation of individual zones, obtaining sufficient quantities of high quality RNA from the elongation zone proved challenging because cells in this region are highly elongated (Supp. Fig. 2). To overcome this limitation, RNA-seq of wheat roots was previously performed using bulk tissue comprising all three zones combined (20). Pooling the zones increased RNA read counts, thereby improving the statistical power for detecting DEGs. Genes identified through both methods and species makes them strong candidates for root angle regulation in both species.

The root cap is the site where gravity perception occurs, because amyloplasts act as statoliths in the columella cells. In response to gravity, these organelles sediment within the cytoplasm and intiate downstream signalling cascade (4). In our experimental setup, root samples were taken from plant grown under stable condition and were not subjected to changes in gravity direction, therefore DEGs are unlikely to be associated with statolith movement but rather involved in the downstream gravitropic or AGO signalling pathways.

In Arabidopsis lateral roots, *CKXs* have been implicated in GSA regulation (11), and two *HvCKXs* genes, *HvCKX1* and *HvCKX5*, were differentially expressed in the barley root cap (Supp. Fig. 3, 4). *HvCKX1* showed its highest expression in seminal roots with medium GSAs, whereas *HvCKX5* showed its lowest expression in the same root class. A third *CKX* gene expressed in the root tissue, *HvCKX3*, showed expression patterns that changed progressively with GSA in the elongation zone. In wheat, orthologues of *HvCKX1, HvCKX5* and *HvCKX3* also showed expressin patterns associated with distinct seminal root classes and their respective GSAs (Fig. 3). These findings suggest that GSA associated regulation of *CKX* gene expression is conserved in cereal seminal roots.

*Hvckx1* mutants showed steeper root systems, stronger gravitropic response and elevated levels of iP-type cytokinins (Fig. 4, Supp. Fig 4). Similarly, exogenous cytokinin application induced steeper root growth in both barley and wheat. Together with the expression of *HvCKX1* in the root cap, these observations suggests that cytokinin levels or the homeostasis of different cytokinin types mediated by *HvCKX1*, may contribute to buffering gravity signalling process downstream of gravity perception in the columella cells. Such a mechanism could dampen gravitropic responsiveness and thereby influence the maintenance of root specific GSAs.

Similar to *Hvckx1, Hvckx3* mutants also displayed steeper root growth angles (Fig. 4). Interestingly, we previously showed that *HvCKX3* expression is upregulated in the elongation zone 12 h after gravistimulation (24). This finding suggests that *HvCKX3* may function in later stages of gravity signalling, by maintaining or re-stablishing the balance between gravitropic response and AGO. In the absence of this regulatory function, *Hvckx3* mutants could have a disturbed balance between these two processes and consequently steeper GSAs. The lower amount of cZ in *Hvckx3* mutants further suggests that this process may involve cytokinin homeostasis within the elongation zone during gravitropic responses. In wheat, AGO is the dominant quantitative regulator of angle determination in seminal root classes (20). Given that *CKX3* orthologues are also expressed in a root class-specific manner in wheat, it is possible that the role of CKX3 in maintaining the between gravitropic response and AGO in seminal root classes is conserved between both species.

In contrast to the steeper root angles observed in barley and wheat seminal roots, both *AtCKX2* mutants and cytokinin treatments result in shallower angles in Arabidopsis lateral roots (11). This contrasting response highlights context-dependent nature of cytokinin signalling during root development and indicates that hormonal functions cannot necessarily be extrapolated across species or root types.

In this study, we showed that barley seminal roots with different GSAs possess distinct transcriptomic profiles, particularly in the root cap. Our approach provides a framework for identifying genes involved in GSA regulation, as exemplified by the *CKX* gene family. However, our analysis did not distinguish between the upper (away from gravity) and lower (towards gravity) sides of the root. Although mutations in *CKX* genes and cytokinin application treatments affected root growth independently of this spatial context, asymmetries relative to the gravity vector are known to be important for establishing GSAs. For example, emerging Arabidopsis lateral roots exhibit asymmetric expression of the auxin efflux carrier PIN3, accompanied by asymmetric auxin signalling (10). Future studies should therefore incorporate spatial information when analysing transcriptomic profiles to further resolve the molecular mechanisms underlying GSA regulation.

Together, our findings identify cytokinin metabolism as a key component of the mechanism that differentiates the GSAs of individual seminal roots. Rather than responding uniformly to the same gravity vector, roots within a single cereal root system maintain distinct growth trajectories, indicating that gravitropic responsiveness is tuned in a root-specific manner. Differential *CKX* expression and cytokinin homeostasis provide a mechanistic basis for modulating this responsiveness and thereby establishing distinct GSAs. The conservation of cytokinin-dependent angle regulation in barley and wheat further suggests that such hormonal tuning is a shared feature of cereal root system architecture. More broadly, our findings show how differential sensitivity to the same directional cue can diversify the growth trajectories of genetically identical organs and thereby contribute to the formation of complex root system architecture.

## Material and methods

### Plant growth material

For barley, the cultivar (cv.) Morex was used for transcriptome analysis and rotation analyses. *ckx* mutants were obtained from Sebastian Gasparis, Plant Breeding and Acclimatization Institute–National Research Institute, Poland (22) in the background cv. Golden Promise. Mutations are described in Suppl. Table 4. For wheat experiments, the hexaploid wheat *Triticum aestivum* cv. Cadenza was used.

### Plant growth conditions

Barley seeds were germinated on moist filter paper at room temperature overnight in darkness. Germinated seeds were transferred to Steel Blue Germination Blotter Paper (Anchor Paper) in A4 observation chambers, wrapped in aluminium foil to maintain roots in darkness (25). Chambers were kept at an 80° angle from vertical in growth incubator, the bottom of the Germination Paper immersed in tap water. The plants were grown in growth cabinets (Conviron) at 18 °C at night (8 h) and 22 °C during the day (16 h). For wheat, similar sized seeds (width = 3.7 ± 0.1 mm, length = 7 ± 0.5 mm) were germinated overnight in the dark at 18 °C on moist germination paper within square petri dishes. Germinated seedlings were transferred to growth pouches made from folded germination paper (Anchor Paper Co., St. Paul, MN, USA) placed in boxes containing autoclaved water. Plants were grown under controlled conditions at 21 °C day/18 °C night temperatures with a 16-h photoperiod.

### Cytokinin treatments

For cytokinin treatments, a 6-BA stock was made in DMSO at a concentration of 100 mM. For barley treatments, 6-BA was added to half-strength Murashige and Skoog medium with MES monohydrate at 0.5g/L final pH of 5.8 and 7g/L agar in the described concentrations and DMSO in the same concentrations as mock treatment. The plants were then transplanted to the medium in A4 observation chambers (25) after germination and grown as described above. For wheat treatments, 6-BA was diluted in water to achieve the desired concentrations. Germinated seedlings in growth pouches were grown in water at the described concentrations.

### Rotation experiments

For gravity-reaction tests, barley seedlings were grown in observation chambers as described and rotated once 45°. Pictures were taken after 0, 3, 6, 9, 12, 24, 48 and 72 h.

### Root phenotyping

Pictures of seedlings on Germination Paper were taken by scanning using a flatbed scanner at 300 dpi. Pictures were analysed in the RootNav software (26). For the rotation experiments, root tip angles were manually measured in ImageJ (27) and the same roots were compared over time.

For the soil-based assays for X-ray µCT, 82-mm-diameter plastic cylinders were uniformly filled with sieved (<2 mm) air-dried loamy sand soil at approximately 1.3 g cm^-3^ bulk density. Soil was collected from the University of Nottingham farm site at Bunny, Nottinghamshire (52°51′41″N, 001°07′42″W). Columns were saturated overnight with water before draining for two days to achieve nominal field capacity. Germinated seedlings were transplanted vertically into the column and covered with 1 cm of loamy sand soil. Three seedlings were planted per column at equal spacing between the centre and edge of the cylinder. Plants were grown for 10 days under the conditions described above. Columns were scanned at the Hounsfield Facility, University of Nottingham, UK, using a Phoenix v|tome|x M 240 kV X-ray computed tomography scanner (Waygate Technologies). Scans were acquired at 50 μm spatial resolution using an X-ray tube voltage of 180 kV voltage and 220 μA current settings. Samples were rotated through 360° in 0.12° increments, generating 3,000 projection images per scan. Each projection was the average of five frames with 333 ms exposure time to reduce image noise. Beam hardening and detector saturation were minimised using a 0.1 mm Cu + 1.0 mm Al filter mounted on the X-ray source. Roots were manually annotated as polylines in VGSTUDIO MAX 2023.1 (Volume Graphics GmbH, Germany), and convex hull volume trait was quantified using an inhouse developed software application for the whole root system.

### Laser Capture Dissection Microscopy (LCM)

For LCM, barley roots with vertical GSA (85-90° from horizontal) with a medium GSA (60-70° from horizontal); and with a shallow GSA (40-50° from horizontal) were sampled. LCM of barley root tips was performed as described previously separating the root cap, meristem and elongation zones (18). Sections of 7-10 roots were pooled per biological replicate.

### RNA-seq analysis barley

RNA-seq reads were mapped against the Morex Transcriptome Atlas (https://ics.hutton.ac.uk/morexgeneatlas/index.html) using Salmon v1.10.3 with default parameters (28, 29). Samples with inconsistent expression of housekeeping genes across the samples were excluded from the analysis (Supplementary Figure 2B, Supplementary table 1). For differential expression analysis, the 3D RNA-seq App was used (30), with the following parameters: Low expressed transcripts and genes were filtered based on analyzing the data mean-variance trend. The expected decreasing trend between data mean and variance was observed when expressed transcripts were determined as those which had ≥ 4 of the 45 samples with count per million reads (CPM) ≥ 1. A gene was expressed if any of its transcripts with the above criteria were expressed. The TMM method was used to normalize the gene and transcript read counts to *log*_2_-CPM. Sample information and raw/expressed gene and transcript number are found in Supplementary Table 1. The limma R package was used for 3D expression comparison, with limma Voom-Weights as Mean Variance Method. For differentially expressed (DE) genes/transcripts, the *log*_2_ fold change (*L*_2_*FC*) of gene/transcript abundance were calculated based on contrast groups (Supplementary table 1) and significance of expression changes were determined using t-test. P-values of multiple testing were adjusted using Benjamini-Hochberg to control the false discovery rate (FDR). A gene/transcript was significantly DE in a contrast group if it had adjusted p-value < 0.05 and *L*_2_*FC* ≥ 1. The related transcript ID in Morex v3 was identified by BLASTn (31). The expression heatmap was created in the R package pheatmap (32) and displays the mean of the TPMs across the biological replicates and the z-score per row. GO term analysis was performed using the agriGO platform (33).

### Phylogenetic analysis

Identifiers and protein sequences are listed in supplementary table 3. Phylogenetic analysis was performed as described in (34), following the pipeline described by (35). Seven cytokinin oxidase/dehydrogenase (*CKX1–CKX7*) genes from *Arabidopsis thaliana* were used as reference sequences. These proteins were characterised by the presence of four conserved domains: IPR006094, IPR015345, IPR016166 and IPR050432. The Arabidopsis CKX protein sequences obtained from The Arabidopsis Information Resource (TAIR) were used to identify putative orthologous CKX candidates in barley and wheat using blastp in BLAST+ (36). Candidate CKX proteins were identified based on sequence similarity and the presence of the conserved CKX domains. Candidates were retained when reciprocal/bidirectional similarity searches supported their orthology to the Arabidopsis CKXs and when they contained the characteristic CKX domain architecture. Protein sequences of the identified CKX candidates were subsequently aligned using MAFFT (37). Alignment quality was assessed and poorly aligned regions and gaps were removed using trimAl (38). The resulting trimmed alignment was used for phylogenetic reconstruction using IQ-TREE (39). Maximum-likelihood phylogenetic trees were generated with 1,000 bootstrap replicates to assess the robustness of the inferred relationships. The resulting phylogenetic tree was exported in Newick format and imported into the Interactive Tree of Life (iTOL) platform (40) for visualisation and annotation. The final tree was presented as an unrooted phylogenetic tree to illustrate the relationships among CKX proteins from Arabidopsis, barley and wheat.

### Auxin and cytokinin metabolite profiling

For hormone metabolite profiling, 1 cm of root tips were sampled 4 days after germination, 5-8 roots per biological replicate, resulting in fresh weight of 4.4 mg – 17 mg (Supplementary Table 5). Samples were collected in liquid nitrogen and freeze-dried overnight. Quantification of cytokinin metabolites was performed according to the method described by (41), including modifications described by (42). Samples (10 mg fresh weight) were homogenized and extracted in 1 ml of modified Bieleski buffer (60% MeOH, 10% HCOOH and 30% H_2_O) together with a cocktail of stable isotope-labelled internal standards (0.2 pmol of CK bases, ribosides, *N*-glucosides, and 0.5 pmol of CK *O*-glucosides, nucleotides per sample added). The extracts were applied onto an Oasis MCX column (30 mg/1 ml, Waters) conditioned with 1 ml each of 100% MeOH and H_2_O, equilibrated sequentially with 1ml of 50% (v/v) nitric acid, 1 ml of H_2_O, and 1 ml of 1M HCOOH, and washed with 1 ml of 1M HCOOH and 1 ml 100% MeOH. Analytes were then eluted by two-step elution using 1 ml of 0.35M NH_4_OH aqueous solution and 2 ml of 0.35M NH_4_OH in 60% (v/v) MeOH solution. The eluates were then evaporated to dryness *in vacuo* and stored at -20°C. Cytokinin levels were determined by an ultra-high performance liquid chromatography-electrospray tandem mass spectrometry (UHPLC-MS/MS) using stable isotope-labelled internal standards as a reference (43). Separation was performed on an Acquity UPLC® i-Class System (Waters, Milford, MA, USA) equipped with an Acquity UPLC BEH Shield RP18 column (150x2.1 mm, 1.7 μm; Waters), and the effluent was introduced into the electrospray ion source of a triple quadrupole mass spectrometer Xevo™ TQ-S MS (Waters). Extraction and purification were performed following the methodology described by (44). Briefly, samples were extracted in cold 50 mmol/L phosphate buffer (pH 7.0) containing 0.1% sodium diethyldithiocarbamate and mixture of stable isotope-labelled internal standards. After the centrifugation (25 200 g, 15 min, 4 °C), a 200 µL aliquot of the supernatant was acidified to pH 2.7 with HCl and purified by in-tip micro solid-phase extraction (in-tip µSPE) using self-assembled columns. After elution, samples were evaporated under reduced pressure, resuspended in 10% aqueous methanol and analyzed by HPLC-MS (1260 Infinity II HPLC system, Agilent Technologies, USA; Kinetex C18 column (50 mmx2.1 mm, 1.7 um; Phenomenex; 6495 Triple Quad detector, Agilent Technologies, USA).

### Statistical analyses

Statistical analyses for the growth experiments and hormone analysis were conducted in R. Treated plants or mutants were compared with untreated controls or WT, respectively, by two-sided Student’s t-tests. One-way ANOVA and Tukey tests were used for multiple comparisons. Differences were considered significant at p < 0.05 marked by * and p <0.01 marked by **.

## Data Availability

RNA-seq data have been deposited in the European Nucleotide Archive (ENA) (https://www.ebi.ac.uk/ena/browser/view/PRJEB123307).

## Acknowledgements

The authors give sincere thanks to Hana Martínková and Andrea Novotná for their help with the phytohormone analyses; to Sebastian Gasparis for sharing the gene-edited barley lines; to Runxuan Zhang and John Brown for advice on the RNA-seq; to Ian Newton and Paul Appleton for help with cryo-sectioning and laser-capture microdissection; and to D.U. Sty and G. Inger for support throughout the project. This work was supported by the Root2Res project (supported by Horizon Europe research and innovation programme under the Grant Agreement no 101060124 and by Innovate UK through the Horizon Europe Guarantee scheme Grant Agreement No. 10106012) and by the Horizon Europe Guarantee MSCA Postdoctoral Fellowship EP/Y014820/1 (GKK); by the BBSRC Discovery Fellowship (BB/S011102/1), the BBSRC New Investigator Research Grant (BB/X014843/1), and the BBSRC Partnering Award (BB/X018806/1) (RB, AB); by the project TowArds Next GENeration Crops, reg. no. CZ.02.01.01/00/22_008/0004581 of the ERDF Programme Johannes Amos Comenius, and the ERC Synergy project “Unravelling Spatio-temporal Auxin Intracellular Redistribution for Morphogenesis” (STARMORPH, reg. no. 101166880) (JKV, ON, AP and PA); by the Scottish Government Strategic Research Programme (2022–2027) under project JHI-D3-1, Healthy Soils for a Green Recovery (TAV); by the Scottish Government’s Rural and Environmental Science and Analytical Services (RESAS) strategic research programme under BARGAIN (JHI-B1-2) (MS); and by the Deutsche Forschungsgemeinschaft (DFG) (499026372 and 470007283 and CIBSS – EXC-2189; 390939984) (JKV, SZF). We thank Research Computing at the James Hutton Institute for providing computational resources and technical support for the “UK’s Crop Diversity Bioinformatics HPC” (BBSRC grants BB/S019669/1 and BB/X019683/1).

## Notes

### Competing Interest Statement

The authors have declared no competing interest.

## References

1. J. P. Lynch, Root phenotypes for improved nutrient capture: an underexploited opportunity for global agriculture. New Phytol. 223, 548–564 (2019).

2. Y. Uga, et al., Control of root system architecture by DEEPER ROOTING 1 increases rice yield under drought conditions. Nat. Genet. 45, 1097–102 (2013).

3. J. Digby, R. D. Firn, The gravitropic set-point angle (GSA): the identification of an important developmentally controlled variable governing plant architecture. Plant. Cell Environ. 18, 1434–1440 (1995).

4. J. Chen, et al., Amyloplast sedimentation repolarizes LAZYs to achieve gravity sensing in plants. Cell 186, 4788-4802.e15 (2023).

5. T. Nishimura, et al., Cell polarity linked to gravity sensing is generated by LZY translocation from statoliths to the plasma membrane. Science (80-.). 381, 1006–1010 (2023).

6. R. Swarup, et al., Root gravitropism requires lateral root cap and epidermal cells for transport and response to a mobile auxin signal. Nat. Cell Biol. 7, 1057–1065 (2005).

7. E. Ghafouri, et al., Article ARF19 acts as a transient auxin response enhancer during root gravitropism ll ARF19 acts as a transient auxin response enhancer during root gravitropism.Cell Rep. 45 (2026).

8. G. B. Monshausen, N. D. Miller, A. S. Murphy, S. Gilroy, Dynamics of auxin-dependent Ca 2+ and pH signaling in root growth revealed by integrating high-resolution imaging with automated computer vision-based analysis. Plant J. 65, 309–318 (2011).

9. W. A. Peer, Y. Cheng, A. S. Murphy, Evidence of oxidative attenuation of auxin signalling. J. Exp. Bot. 64, 2629–2639 (2013).

10. M. R. Rosquete, et al., An auxin transport mechanism restricts positive orthogravitropism in lateral roots. Curr. Biol. 23, 817–822 (2013).

11. S. Waidmann, et al., Cytokinin functions as an asymmetric and anti-gravitropic signal in lateral roots. Nat. Commun. 10, 3540 (2019).

12. S. Z. Farkas, et al., SHOOT GRAVITROPISM 9 links sensory timing to the initial lateral root growth angle. Proc. Natl. Acad. Sci. U. S. A. 123, 1–8 (2026).

13. S. Roychoudhry, et al., Antigravitropic PIN polarization maintains non-vertical growth in lateral roots. Nat. Plants 9, 1500–1513 (2023).

14. F. Hochholdinger, W. J. Park, M. Sauer, K. Woll, From weeds to crops: genetic analysis of root development in cereals. Trends Plant Sci. 9, 42–48 (2004).

15. X. Kong, et al., Ethylene regulates auxin-mediated root gravitropic machinery and controls root angle in cereal crops. Plant Physiol. 195, 1969–1980 (2024).

16. Y. Xiong, et al., ABA-auxin cascade regulates crop root angle in response to drought. Curr. Biol. 35, 542-553.e4 (2025).

17. X. Song, et al., Auxin controls rice root angle via kinase OsILA1-mediated cell wall modifications. Sci. Adv. 11, eady2320 (2025).

18. G. K. Kirschner, et al., ENHANCED GRAVITROPISM 2 encodes a STERILE ALPHA MOTIF–containing protein that controls root growth angle in barley and wheat. Proc. Natl. Acad. Sci. 118, e2101526118 (2021).

19. R. Fusi, et al., Root angle is controlled by EGT1 in cereal crops employing an antigravitropic mechanism. Proc. Natl. Acad. Sci. 119, 1–10 (2022).

20. A. Bansod, et al., Growth angles of distinct root classes in wheat are determined by a gradient of anti-gravitropic activity. Res. Sq. 1–22 (2026).

21. K. Mrízová, et al., Overexpression of Cytokinin Dehydrogenase Genes in Barley (Hordeum vulgare cv. Golden Promise) Fundamentally Affects Morphology and Fertility. PLoS One 8, e79029 (2013).

22. S. Gasparis, M. Przyborowski, M. Kała, A. Nadolska-Orczyk, Knockout of the HvCKX1 or HvCKX3 Gene in Barley (Hordeum vulgare L .) by RNA-Guided Cas9 Nuclease Root Morphology. Cells 8 (2019).

23. F. Stemp-Walsh, et al., Roots of wheat and rice maintain gravitropic setpoint angles. J. Exp. Bot. 77, 1–9 (2026).

24. L. Guo, et al., ENHANCED GRAVITROPISM coordinates molecular adaptations to gravistimulation in the elongation zone of barley roots. New Phytol. 0–2 (2023). 10.1111/nph.18717.

25. A. G. Bengough, et al., Gel observation chamber for rapid screening of root traits in cereal seedlings. Plant Soil 262, 63–70 (2004).

26. R. Yasrab, et al., RootNav 2.0: Deep learning for automatic navigation of complex plant root architectures. Gigascience 8, 1–16 (2019).

27. J. Schindelin, et al., Fiji: an open-source platform for biological-image analysis. Nat. Methods 9, 676–682 (2012).

28. W. Guo, et al., A barley pan-transcriptome reveals layers of genotype-dependent transcriptional complexity. Nat. Genet. 57, 441–450 (2025).

29. R. Patro, G. Duggal, M. I. Love, R. A. Irizarry, C. Kingsford, Salmon provides fast and bias-aware quantification of transcript expression. Nat. Methods 14 (2017).

30. W. Guo, et al., 3D RNA-seq: a powerful and flexible tool for rapid and accurate differential expression and alternative splicing analysis of RNA-seq data for biologists. RNA Biol. 18, 1574–1587 (2021).

31. M. Mascher, et al., Long-read sequence assembly: A technical evaluation in barley. Plant Cell 33, 1888–1906 (2021).

32. R. Kolde, pheatmap: Pretty Heatmaps R package version 1.0.13. (2025).

33. T. Tian, et al., AgriGO v2.0: A GO analysis toolkit for the agricultural community, 2017 update. Nucleic Acids Res. 45, W122–W129 (2017).

34. R. Fusi, et al., The auxin efflux carrier PIN1a regulates vascular patterning in cereal roots. New Phytol. 244, 104–115 (2024).

35. S. K. Mutte, D. Weijers, High-resolution and Deep Phylogenetic Reconstruction of Ancestral States from Large Transcriptomic Data Sets. Bio-protocol 10, 1–14 (2020).

36. C. Camacho, et al., BLAST + : architecture and applications. BMC Bioinformatics 10, 1–9 (2009).

37. K. Katoh, K. Misawa, K. Kuma, T. Miyata, MAFFT : a novel method for rapid multiple sequence alignment based on fast Fourier transform. Nucleic Acids Res. 30, 3059–3066 (2002).

38. S. Capella-Gutiérrez, J.M. Silla-Martínez, T. Gabaldón, trimAl: A tool for automated alignment trimming in large-scale phylogenetic analyses. Bioinformatics 25, 1972–1973 (2009).

39. L. Nguyen, H. A. Schmidt, A. Von Haeseler, B. Q. Minh, IQ-TREE : A Fast and Effective Stochastic Algorithm for Estimating Maximum-Likelihood Phylogenies. Mol. Biol. Evol. 32, 268–274 (2014).

40. I. Letunic, P. Bork, Interactive Tree Of Life (iTOL) v5 : an online tool for phylogenetic tree display and annotation. Nucleic 49, 293–296 (2021).

41. J. Svačinová, et al., A new approach for cytokinin isolation from Arabidopsis tissues using miniaturized purification: pipette tip solid-phase extraction. Plant Methods 8, 17 (2012).

42. I. Antoniadi, et al., Cell-Type-Specific Cytokinin Distribution within the Arabidopsis Primary Root Apex. Plant Cell 27, 1955–1967 (2015).

43. D. Rittenberg, G. L. Foster, A new procedure for quantitative analysis by isotope dilution, with application to the determination of amino acids and fatty acids. J. Biol. Chem. 133, 737–744 (1940).

44. P. Hladík, I. Petřík, A. Žukauskaitė, O. Novák, A. Pěnčík, Metabolic profiles of 2-oxindole-3-acetyl-amino acid conjugates differ in various plant species. Front. Plant Sci. 14, 1–13 (2023).

