## Supplemental Figures for "Cytokinin regulates the gravitropic setpoint angle of seminal roots in barley and wheat"

**Supplementary Figures Bonsad et. al**

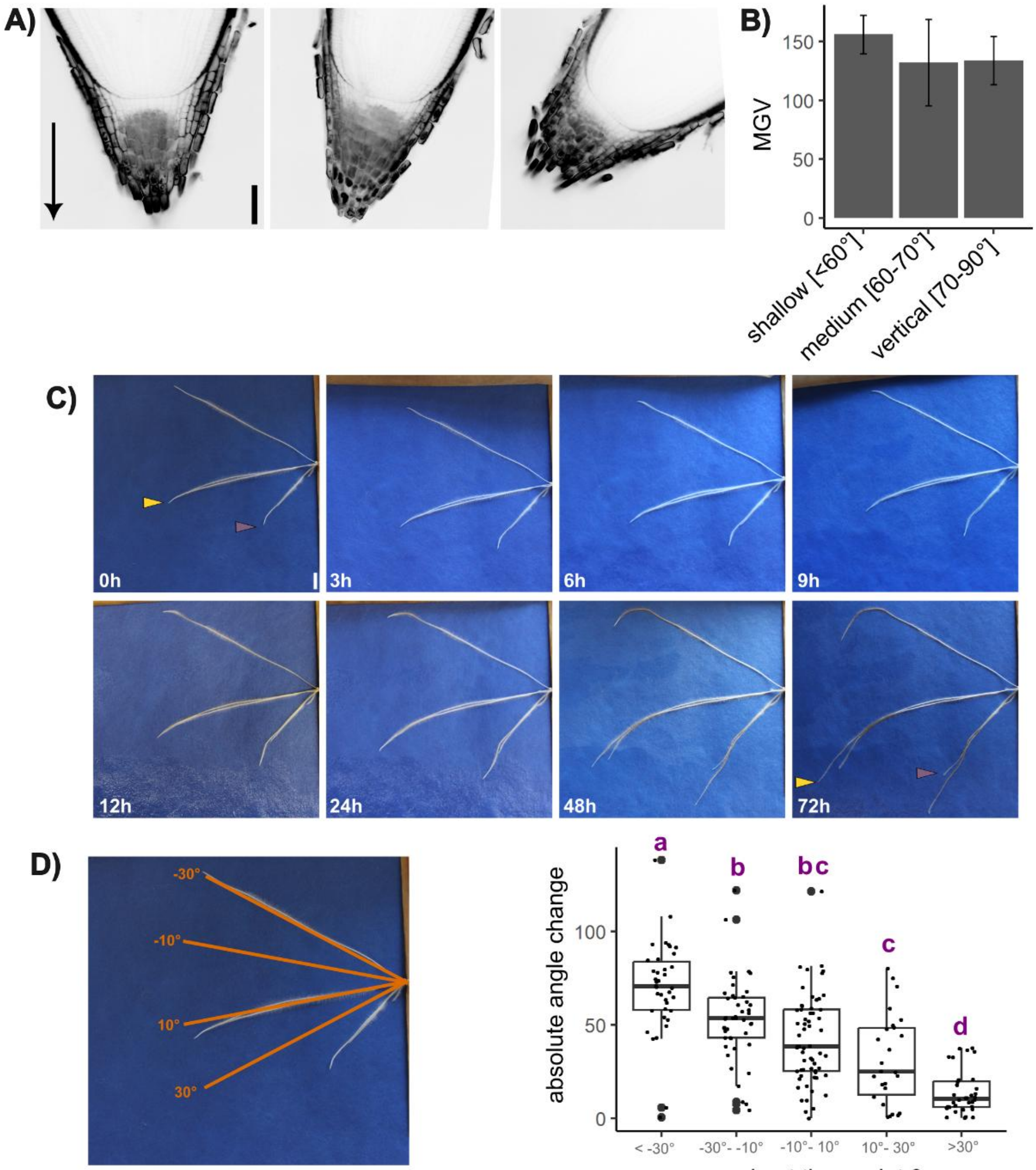


**Supplementary Figure 1: Barley seminal roots have a set angle in relation to gravity (gravitropic set-point angle, GSA).**

A) Barley seminal roots (4 days after germination) with different GSA (left ~86°, middle ~74° right ~45°) show similar starch amount and distribution in their root cap, visualised by modified pseudo-Schiff propidium iodide (mPS-PI) staining; arrowhead shows direction of gravity; scale bar 100 µm.
B) Quantification of starch by measuring the mean gray value (MGV) in the root cap; bar plot shows mean and standard deviation of roots with vertical, medium and shallow angle; a one-way ANOVA and Tukey test showed no significant differences; n= 12 roots.
C) After a gravistimulation by a 90° rotation, barley seminal roots return to their gravitropic set-point angle over time. Roots that are misplaced from their GSA by the rotation (yellow arrowhead; ~ 60° before the rotation) adapt their growth direction to return to the GSA; roots that kept the GSA after the rotation (~45°) (magenta arrowhead) keep their growth direction; scale bar 1 cm.
D) Absolute change in angle of roots after 72 h after a 90° rotation depending on their starting angle. Roots with an angle of -30° or smaller adapt their angle the strongest, while roots with an angle of 30° or more vertical keep their growth direction and adapt their angle less. Treatments that share a common letter are not significantly different (one-way ANOVA and Tukey test); every data point presents one root; n = 38 (< -30°), 45 (-30° – -10°), 61 (-10° – 10°), 26 (10° – 30°), 36 (>30°).

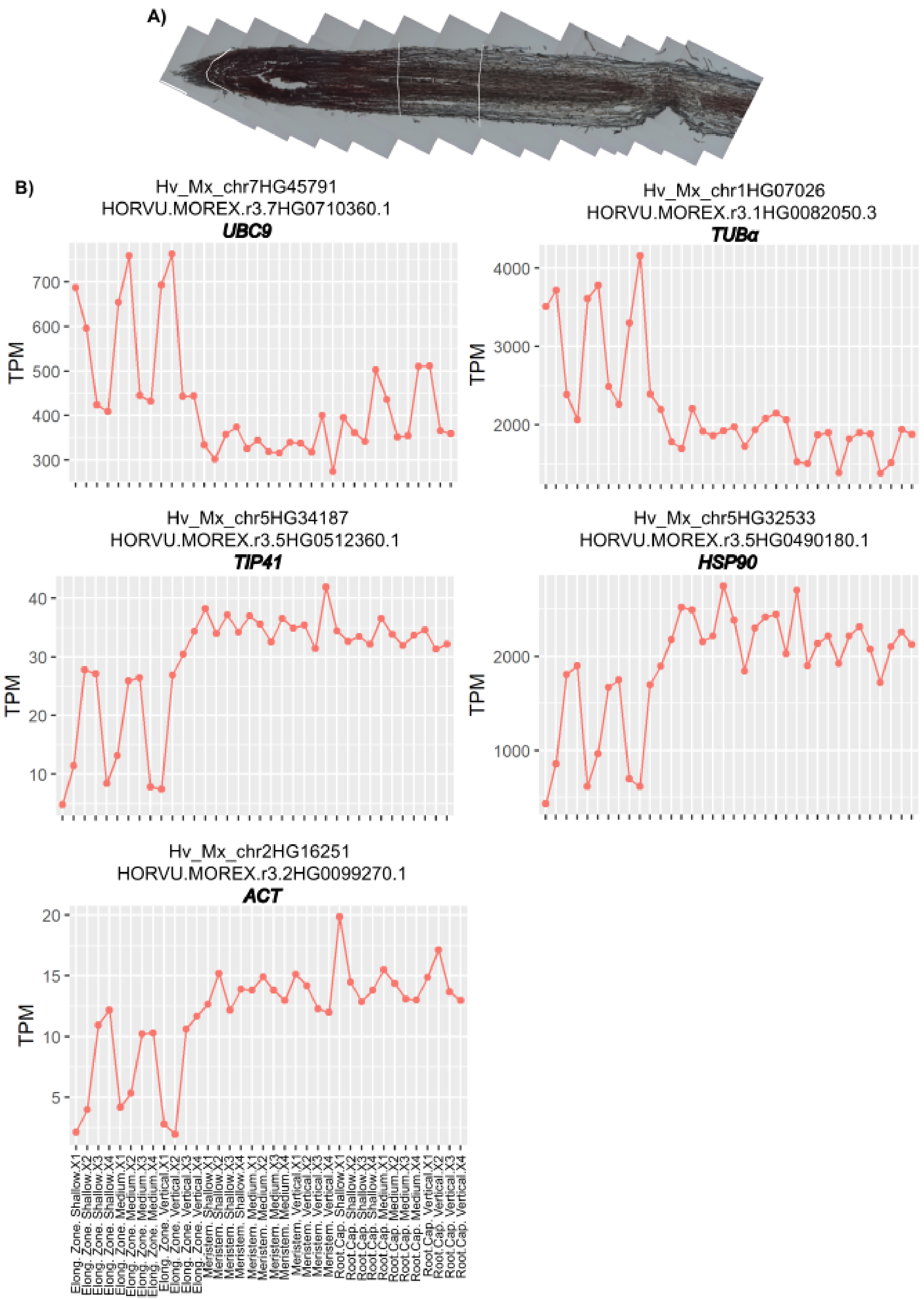


**Supplementary Figure 2: Root zone samples were collected by laser capture microdissection (LCM).**

A) Exemplary picture of root zones sampled at LCM; root cap, meristem, elongation zone were sampled as indicated; scale bar 150 µm.

B) Based on transcript per million values of housekeeping genes, biological replicates with expression divergent from the other samples were removed from further analysis (Elongation Zone. Shallow.X1; Elongation Zone. Shallow.X2; Elongation Zone. Medium.X1; Elongation Zone. Medium.X2; Elongation Zone. Vertical.X1; Elongation Zone. Vertical.X2); ubiquitin-conjugating enzyme 9 (*UBC9*) (1); *ACTIN* (*ACT*); *TUBULINα* (*TUBα*) (2); TIP41-like family protein (*TIP41*) (1); heat shock protein (*HSP90*) (2).

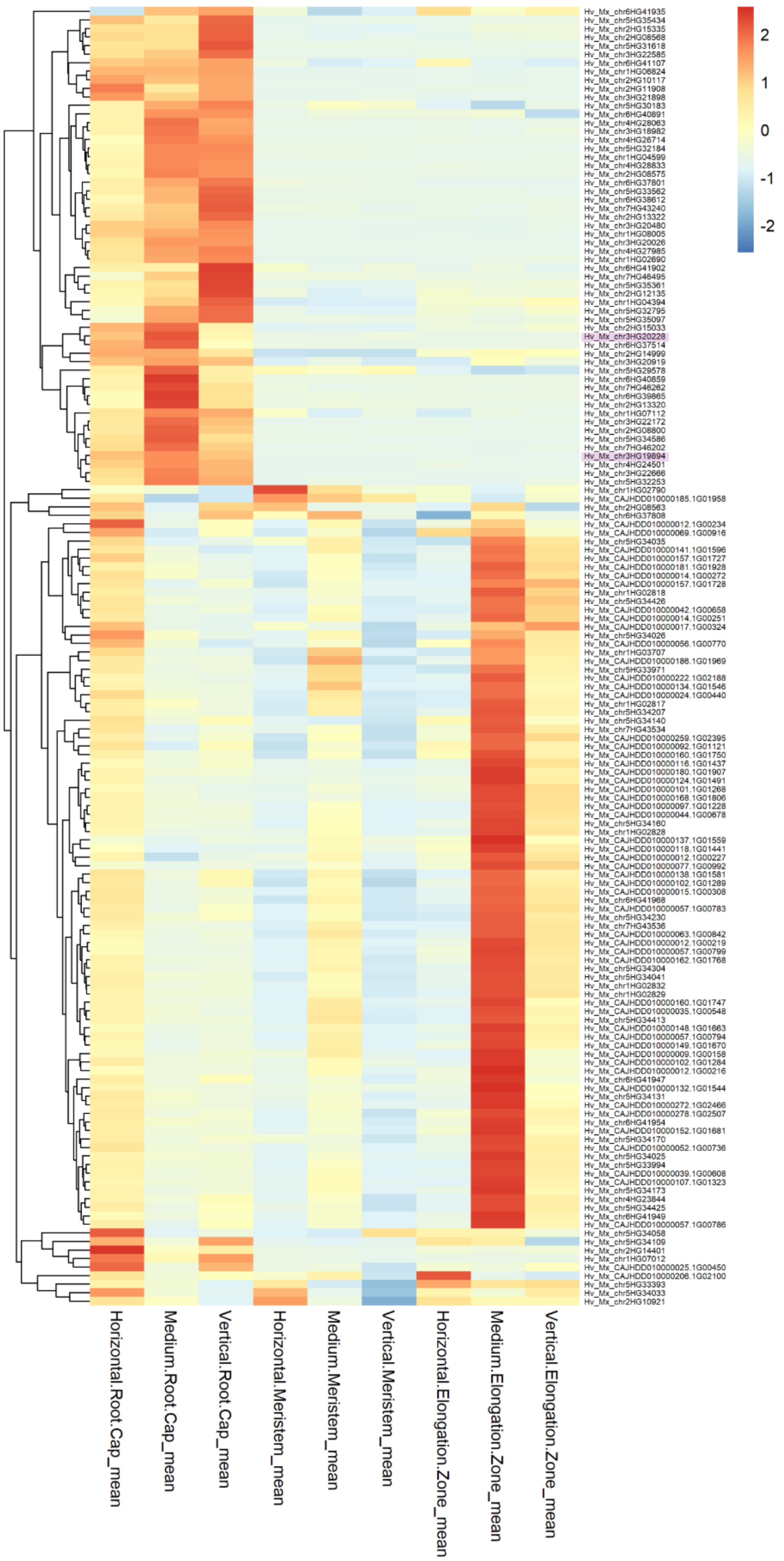


**Supplementary Figure 3: Heatmap of differentially expressed genes in the barley root tip.** *HvCKXs* are highlighted in purple; colours show z-score according to colour scale.

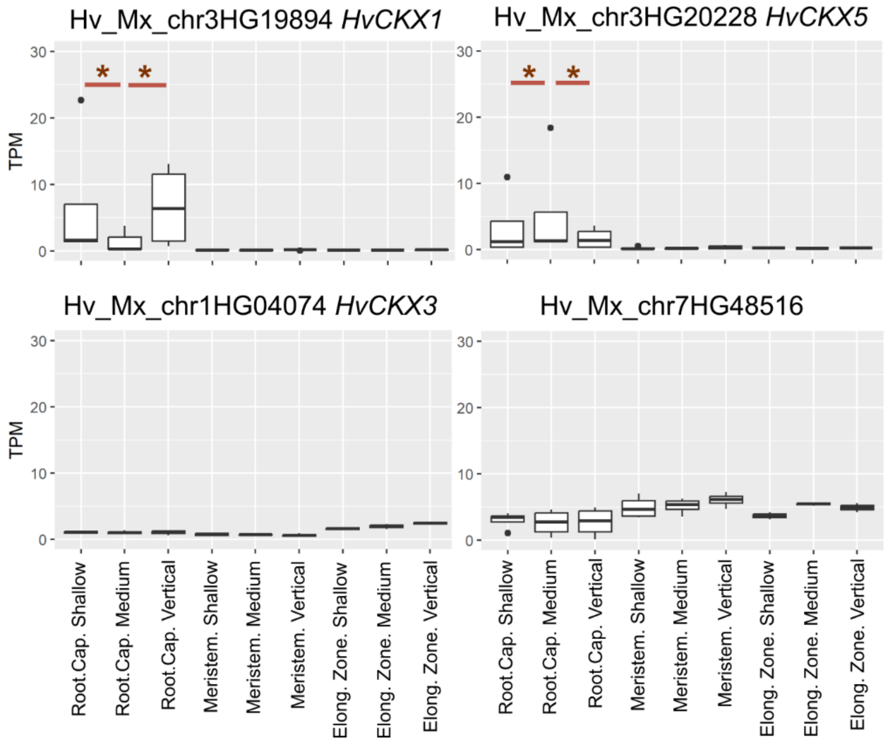


**Supplementary Figure 4: Expression of *HvCKXs* in the barley root tip.**
Expression of all root-expressed *HvCKXs* in the root cap, meristem and elongation zone in roots with different angles; TPM = transcript per million; samples that are significantly differentially expressed in a comparison of roots with different angles within each zone are marked by asterisks.

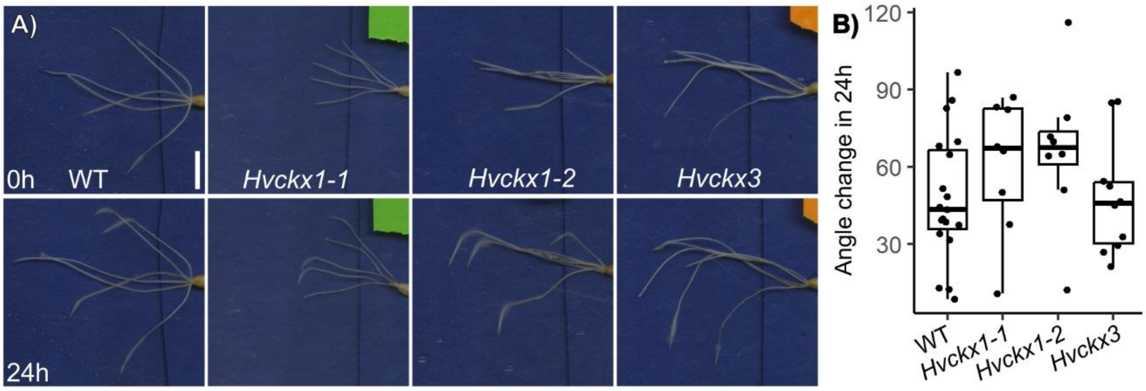


**Supplementary Figure 5: Gravity response of *Hvckx* mutants**
A) *Hvckx* mutants have a faster gravity response. WT, *Hvckx1-1*, *Hvckx1-2* and *Hvckx3* before (0 h) and 24 h after a 90° rotation; scale bar 1 cm.
B) Root angle change in relation to start angle 24 h after rotation; n = 19 roots (WT), 8 (*Hvckx3*), 8 (*Hvckx1-1*), 12 (*Hvckx1-2*).

**Supplementary Video 1:** X-ray microcomputed tomography (µCT) video of wild-type barley cv. Golden promise grown in soil columns for 10 days after germination. Snapshot shown in Figure 4.

**Supplementary Video 2:** X-ray microcomputed tomography (µCT) video of barley mutant *Hvckx1-1* grown in soil columns for 10 days after germination. Snapshot shown in Figure 4.

**Supplementary Video 3:** X-ray microcomputed tomography (µCT) video of barley mutant *Hvckx1-2* grown in soil columns for 10 days after germination. Snapshot shown in Figure 4.

**Supplementary Video 4:** X-ray microcomputed tomography (µCT) video of barley mutant *Hvckx3* grown in soil columns for 10 days after germination. Snapshot shown in Figure 4.

### References

1. W. Hua, *et al.*, Identification of Suitable Reference Genes for Barley Gene Expression Under Abiotic Stresses and Hormonal Treatments. *Plant Mol. Biol. Report.* **33**, 1002–1012 (2015).

2. J. Cai, *et al.*, Selection of appropriate reference genes for the detection of rhythmic gene expression via quantitative real-time PCR in Tibetan hulless barley. *PLoS One* **13**, 1–19 (2018).
